# Deciphering how interneuron specific 3 cells control oriens lacunosum-moleculare cells to contribute to circuit function

**DOI:** 10.1101/2020.09.21.306472

**Authors:** Alexandre Guet-McCreight, Frances K Skinner

**Author notes:** **Corresponding Author:** Alexandre Guet-McCreight, University of Toronto.

## Abstract

The wide diversity of inhibitory cells across the brain makes them suitable to contribute to network dynamics in specialized fashions. However, the contributions of a particular inhibitory cell type in a behaving animal are challenging to untangle as one needs to both record cellular activities and identify the cell type being recorded. Thus, using computational modeling and theory to predict and hypothesize cell-specific contributions is desirable. Here, we examine potential contributions of interneuron-specific 3 (I-S3) cells - an inhibitory interneuron found in CA1 hippocampus that only targets other inhibitory interneurons - during simulated theta rhythms. We use previously developed multi-compartment models of oriens lacunosum-moleculare (OLM) cells, the main target of I-S3 cells, and explore how I-S3 cell inputs during *in vitro* and *in vivo* scenarios contribute to theta. We find that I-S3 cells suppress OLM cell spiking, rather than engender its spiking via post-inhibitory rebound mechanisms, and contribute to theta frequency spike resonance during simulated *in vivo* scenarios. To elicit recruitment similar to *in vitro* experiments, inclusion of disinhibited pyramidal cell inputs is necessary, implying that I-S3 cell firing broadens the window for pyramidal cell disinhibition. Using *in vivo* virtual networks, we show that I-S3 cells contribute to a sharpening of OLM cell recruitment at theta frequencies. Further, shifting the timing of I-S3 cell spiking due to external modulation shifts the timing of the OLM cell firing and thus disinhibitory windows. We propose a specialized contribution of I-S3 cells to create temporally precise coordination of modulation pathways.

**New & Noteworthy:** How information is processed across different brain structures is an important question that relates to the different functions that the brain performs. Using modelling and theoretical analyses, we show that an inhibitory cell type that only inhibits other inhibitory cells can broaden the window for disinhibition of excitatory cells, manage input pathway switching, and modulate inhibitory cell spiking. This work contributes to knowledge of how coordination between sensory and memory consolidation information can be attained.

## Introduction

Across the brain there is a variety of different types of excitatory and inhibitory neurons that control how information is processed (1–7). This diversity spans morphological, electrophysiological and molecular aspects, and examination of specific inhibitory cell types shows that there are distinct neuronal subclasses that can be mapped to function and behaviour (8). However, identifying a cell type goes beyond characterization of any single genetic marker making such mappings a difficult endeavour. Explorations of inhibitory cells have additional challenges since they are typically smaller in size and are rarer than their excitatory cell counterparts, making them more difficult to record from *in vivo*. Even though there are conceptual and technical challenges in classifying cell types, it is clear, for example, that brain diseases can be specific in the cell types that they affect (9).

The CA1 hippocampus, a brain area associated with memory formation (10), stands out in the field by the vast amount of experimental literature characterizing its inhibitory cell types (2, 4, 6, 11). In particular, there are four known types of interneuron-specific (I-S) cells -inhibitory cells that are specialized to primarily target other interneurons, and not pyramidal (PYR) cells (12–18). Of them, we here focus on one type, the interneuron-specific 3 (I-S3) cell type via consideration of its primary cellular targets. I-S3 cells express vasoactive intestinal polypeptide (VIP) and calretinin (CR), and their primary targets are oriens lacunosum moleculare (OLM) cells (12, 15), a cell type that expresses somatostatin (SOM) and inhibits the distal apical dendrites of PYR cells in CA1 (19, 20). PYR cells represent by far the largest proportion of neurons in the CA1 network (2, 6). Inputs carrying “sensory-related information” from entorhinal cortex synapse onto their distal dendrites, and inputs carrying “retrieval-related information” from CA3 synapse onto their proximal dendrites (4, 21). The OLM cells have been shown to gate sensory encoding via the inhibition of PYR cell distal dendrites (20, 22), and I-S3 cells are thus well placed for input-specific information gating in the CA1 area. This dis-inhibitory circuitry (i.e., I-S3 cell inhibition of OLM cells causing disinhibition of PYR cells) is not necessarily unique to the hippocampus since strikingly similar circuitries with VIP+ and SOM+ cell types have also been reported across several different areas of cortex (17). Generally speaking, activation of VIP+ cells *in vivo* across different cortical areas has been associated with improved performances in a variety of different learning and memory paradigms as well as the facilitation of synaptic potentiation in PYR cells (17).

From *in vitro* studies, it has been shown that I-S3 cells might exert control over OLM cells via post-inhibitory rebound (PIR) mechanisms (15). However, it is possible that inhibitory input from I-S3 cells could suppress OLM cell firing or I-S3 cells may indirectly control OLM cell firing via other cellular pathways. Further, it is unknown how and whether *in vitro* observations would translate to *in vivo* settings, and targeting and dissecting out the contributions of I-S3 cell inputs to OLM cell spiking *in vivo* would be technically difficult and challenging. To begin to understand the inhibitory control that I-S3 cells exert over OLM cells, it is first important to understand the various input integration mechanisms that govern OLM cell dendritic physiology.

In addition to having highly active dendritic properties (23–25), OLM cells exhibit prominent sag currents due to hyperpolarization-activated cation channels (h-channels) (26–28), which can also promote PIR spiking (29). H-channels are also thought to contribute to spike resonance properties in OLM cells *in vivo* at theta frequencies, but when tested *in vitro* using dynamic-clamp experimental techniques, OLM cells phase-locked well to theta frequency-modulated inputs, independently of whether h-channels were blocked (30) . However, since dynamic-clamp experiments are largely limited to somatic injections, possible contributions of dendritic h-channels were not assessed. Previous computational modelling work did explore OLM cell dendritic h-channel contributions to spike resonance in which OLM cell models with synapses distributed onto their somatic and dendritic compartments to generate simplified *in vivo*-like states were used (31). It was found that there is a shift in spike resonance from high to low theta frequencies if, respectively, dendritic h-channels are present or not (31). Subsequently, a “neuron-to-model matched” (NMM) approach was used to create new OLM cell models in which there was a tight integration of experimental data with the multi-compartment model building, so as to determine whether dendritic h-channels are in fact present or absent (27) . These new OLM models created using this NMM approach strongly suggested that h-channels must be present on OLM cell dendrites.

In parallel, in previous work using I-S3 cell models (32), we had developed an approach for performing high-resolution parameter searches to find excitatory and inhibitory synaptic input parameter combinations that could generate *in vivo*-like states, and thus create virtual networks from a particular cell’s perspective. We took advantage of this approach to do the same for these new OLM cell models where we demonstrated that h-channel dendritic currents are consistently enhanced during *in vivo*-levels of synaptic bombardment (33). Importantly, I-S3 cell inputs are a major source of inhibitory synaptic inputs to OLM cells (13) . Overall, these model developments provide an opportunity to do a detailed *in silico* exploration of the potential contributions of I-S3 cells in a behaving animal via its control over OLM cells during theta activities, given all of our previous findings on OLM cell dendritic h-channels (27, 31), advancements in generating detailed *in vivo* virtual network simulations (32, 33), and discovery of the activation patterns of I-S3 cells during animal behaviour (16).

In this study, we use computational modelling and theoretical analyses to examine how I-S3 cells manifest control over OLM cells so as to decipher how these specialized cell types contribute to function. In the strategy implemented in this work, we use the new OLM cell models (27) with synaptic inputs from I-S3 cells as well as other inputs (33), and perform simulations with these models under *in vitro* and *in vivo*-like contexts (see **Fig 1** illustration), as motivated by experimental data specific to OLM and I-S3 cells. Our explorations in the *in vitro* context are based on observations from Tyan et al. (15) which showed that I-S3 cells could preferentially control the timing of OLM cell spiking at 5 and 10 Hz (theta frequencies) but not at non-theta frequencies. Our explorations in the *in vivo* context are based on results from Luo et al. (16) demonstrating that I-S3 cells are activated with a delay relative to theta-run epochs and spike during the rising to peak phases of theta rhythms. Our models allow a focus on biophysical details of OLM cells and we use theoretical phase response curve (PRC) analyses from *in vitro* aspects to motivate and understand mechanisms of entrainment and spiking resonance *in vivo* in considering specialized contributions of I-S3 cells. In the end, our modeling study predicts that I-S3 cells contribute to function by suppressing spiking in OLM cells and broadening the window for the synaptic disinhibition of pyramidal cells. Overall, our work shows that close interfacing of computational and experimental studies can help us make progress toward the challenging endeavour of mapping specialized cell types to function and behaviour.

**Figure 1:**
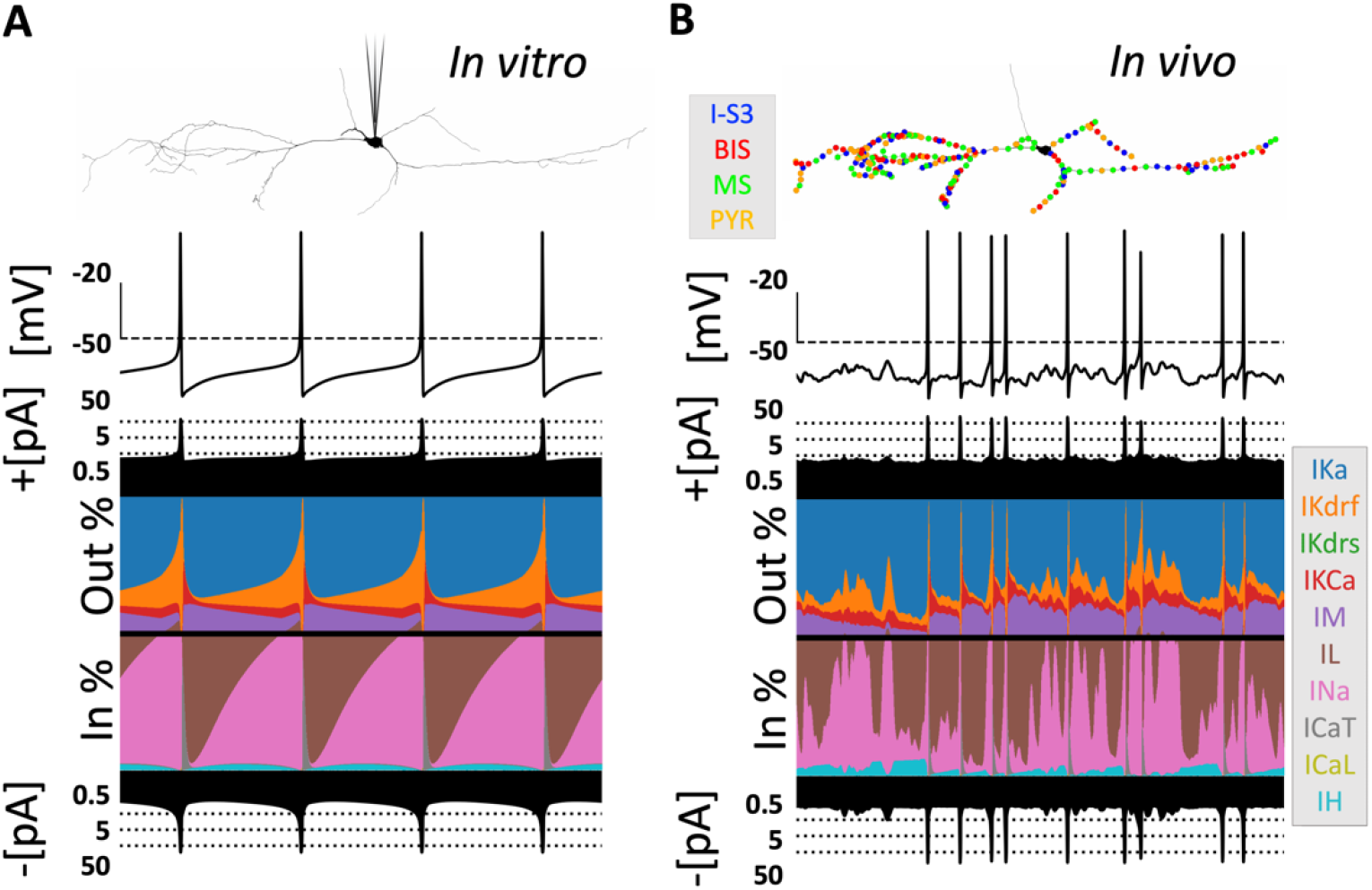
OLM cell model output in *in vitro* and *in vivo* contexts. **A**: Schematic demonstrating the *in vitro* context where DC current is injected directly into the OLM cell model somatic compartment (top) such that spiking is generated. Currentscape plots (74) are shown below, where in each currentscape plot, the top trace is the voltage trace (y-axis scale bar = -50 mV [horizontal dashed line] to -20 mV), the filled-in black traces above and below the coloured plots are the total outward and inward currents respectively (dotted lines = ± 0.5 pA, ± 5 pA, ± 50 pA), and the coloured plots (see colour references in the legend on the right) show the percent contributions of each individual outward (top half of the plot) and inward (bottom half of the plot) current to the total outward and inward currents. B: Schematic demonstrating the *in vivo* context where the OLM cell model dendritic compartments are bombarded with synaptic inputs from different excitatory (PYR) and inhibitory (I-S3, BIS: Bistratified cell inputs, MS: Medial Septal inputs) presynaptic populations (top) such that the voltage dynamics exhibit characteristics that are typical of a high-conductance state. As in panel A, the currentscape plot is shown below.

## Materials and Methods

Associated code for all of the simulations that are presented in this work can be found at the following link: https://github.com/FKSkinnerLab/OLM_IS3control.

### OLM cell models

The models upon which the present work is based have been previously published. Detailed descriptions of these models with equations and parameter values can be accessed from Guet-McCreight and Skinner (33) and Sekulic et al. (27). Here we provide a brief description:

Two detailed multi-compartment OLM cell models, cell 1 and cell 2, were developed in NEURON (34) using a “neuron-to-model matched” (NMM) approach (27). That is, the morphology, passive properties and h-channel characteristics in each model were specific to cell 1 or 2 as measured in the biological OLM cell, and other biophysical currents were optimized using electrophysiological data specific to cell 1 or 2. As expected, model parameter values, such as the biophysical channel conductances, differed between cell 1 and cell 2. Both cell models were constructed using morphological and electrophysiology data from intermediate CA1 mouse hippocampus. Code for the models is available from https://github.com/FKSkinnerLab/OLMng. Ion channel mechanisms include: hyperpolarization-activated cation channels (*H*), transient sodium (*NaT*), fast and slow delayed rectifier potassium (*Kdrf, Kdrs*), A-type potassium (*A*), M-type potassium (*M*), T-and L-type calcium (*CaT, CaL*), and calcium-dependent potassium (*KCa*) channels. All of these mechanisms are distributed throughout the cell, with *CaT, CaL*, and *KCa* being inserted only in dendritic compartments, the rest inserted in somatic and dendritic compartments, and *NaT, Kdrf*, and *Kdrs* also being inserted in axonal compartments. We refer to currents generated by the different ion channel mechanisms with subscripts.

These two OLM cell models developed using an NMM approach are the most up-to-date OLM cell models currently available (27), and optimized fits to the electrophysiological data support the inclusion of h-channels on the dendrites of OLM cells. Although each of these models represent OLM cells as captured by mimicking the electrophysiological recording outputs from the particular biological OLM cell, they differ in their detailed morphologies and conductance values for each of the various ion channel mechanisms. As such, they encapsulate cellular heterogeneity as is known to exist (35). For example, cell 1 has a maximal h-channel conductance of 0.1063 pS/µm^2^ whereas it is 0.0608 pS/µm^2^ in cell 2. All parameter values can be found in Guet-McCreight and Skinner (33) and Sekulic et al. (27). Throughout the paper we use both models so as to consider whether these differences would affect our results.

### Synapse models

For the synapse model we use NEURON’s built-in Exp2Syn function. The input populations to OLM cells include: I-S3 cell inputs, GABAergic long-range projecting inputs from medial septum (MS), bistratified cell inputs (BIS), and local PYR cell inputs [I-S3 cells & MS: see Chamberland et al. (13); BIS & PYR cells: see Leão et al. (20)]. In the absence of specific spatial constraints, these inputs are distributed randomly across all dendritic compartments. OLM cells also receive inputs locally from long-range projecting VIP+ cells in CA1 (18), though because these cells are silent during theta rhythms, we did not include them in this study. In previous work, we had performed optimizations across dendritic compartments so as to estimate the synaptic weights for each input type that matched the experimentally recorded synaptic current amplitudes measured at the soma, also taking into consideration measured synaptic kinetics [see Table 2 in Guet-McCreight and Skinner (33)]. In doing this, we obtained increasing values with distance from soma (cell 1: *G*_*PYR*_ = 0.00020 to 0.00082 µS; *G*_*MS*_ = 0.00024 to 0.00132 µS; *G*_*I*−*S*3_ = 0.00018 to 0.00068 µS; *G*_*BIS*_ = 0.00021 to 0.00100 µS; cell 2: *G*_*PYR*_ = 0.00020 to 0.00120 µS; *G*_*MS*_ = 0.00024 to 0.00132 µS; *G*_*I*−*S*3_ = 0.00018 to 0.00120 µS; *G*_*BIS*_ = 0.00021 to 0.00100 µS). Consideration of other inputs and further details can be found in Guet-McCreight and Skinner (33).

#### In vitro model explorations

##### *In vitro* setup

We use our models to consider an *in vitro* context as used by Tyan et al. (15) of hippocampal slice recordings. In these experiments, although excitatory and inhibitory synapses were not specifically blocked during intracellular OLM cell recordings, cells are relatively silent compared to *in vivo* recordings, and so we do not include any additional synaptic bombardment (see **Fig. 1A**). To model this context, we apply various holding currents to the somatic compartment and rhythmic synaptic input activations on dendritic compartments, which is analogous to the optogenetic experiments performed by Tyan et al. (15). More specifically, we use a similar protocol in our computational OLM cell models where we first depolarize the cell enough (cell 1: *I*_*in j*_ = 49 pA; cell 2: *I*_*in j*_ = 45 pA) to attain a spike rate of 7.25 Hz. We then activate select input populations at different frequencies in our models. The synaptic locations for the inputs are chosen randomly across the dendritic arbour of OLM cells (**Fig. 2**) using our previously developed techniques (33). We look at a variety of different input schemes to gauge possible contributions of different network components, but we mainly focus our explorations on the contributions from I-S3 cell inputs. When PYR cell inputs are included together with inhibitory ones, they are activated with a delay that has been observed experimentally (see section below).

**Figure 2:**
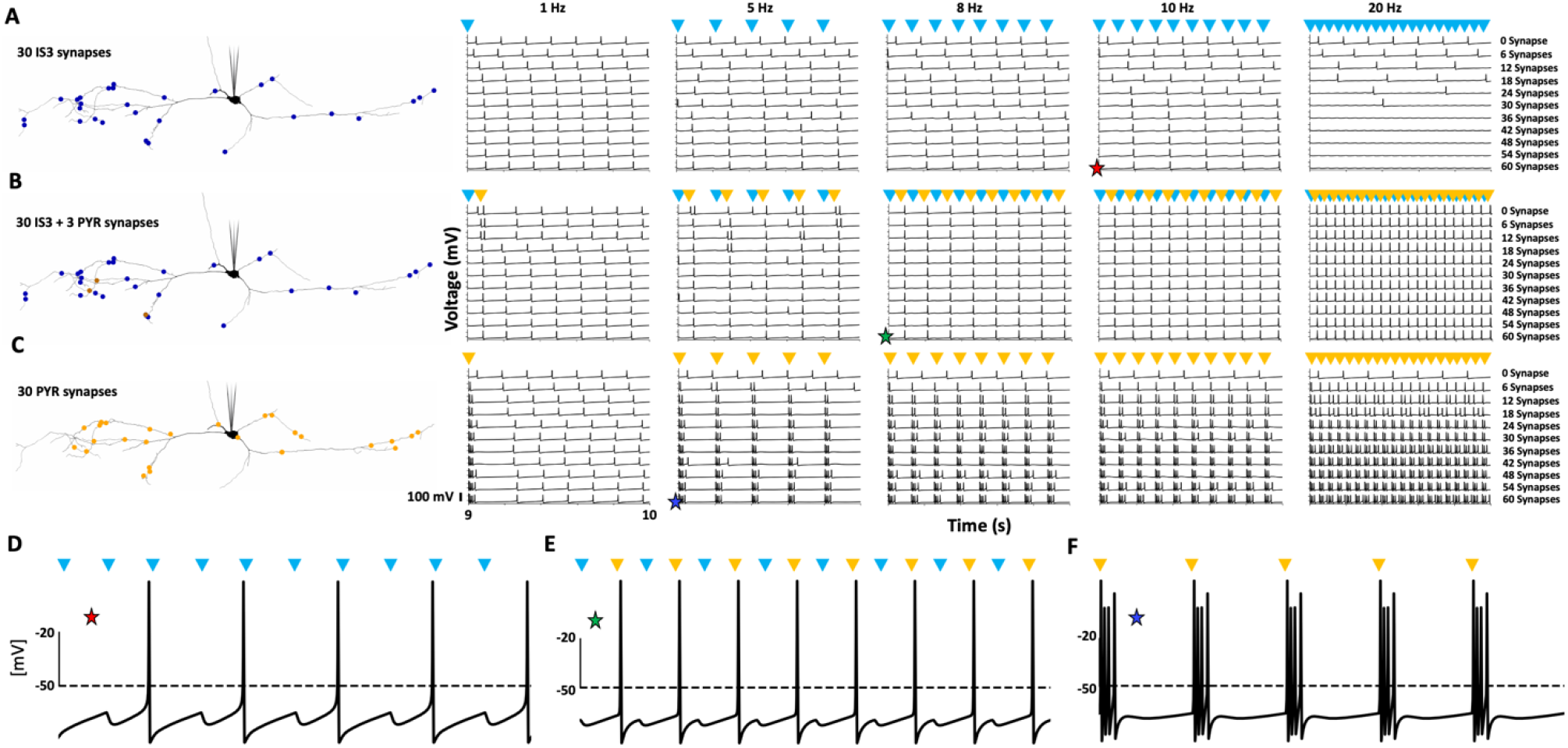
I-S3 cell inputs alone cannot entrain OLM cell spiking at frequencies higher than their baseline spike rate. A: Example synaptic locations (left) of 30 I-S3 cell inputs onto an OLM cell model (note that synaptic locations can be overlayed). Simulated voltage traces (right) in the 9-10s simulation times across several frequency-modulated I-S3 cell input frequencies (left to right) and numbers of I-S3 cell synapses (top to bottom). Blue arrows indicate times when I-S3 cell synapses are activated. For each voltage trace axis, the two notches indicate voltages of -50 mV and 0 mV. B: Example synaptic locations (left) of 30 I-S3 cell inputs and 3 PYR cell inputs onto an OLM cell model. Simulated voltage traces (right; as in A) but with inclusion of 3 PYR cell inputs (timing indicated by the yellow arrows). Note that this number of PYR cell inputs is fixed across simulations. The times at which the 3 PYR cell inputs are activated are indicated by the yellow arrows and start with a delay of 78.125 ms relative to I-S3 cell inputs, consistent with their relative timing during theta rhythms (4, 16). C: Example synaptic locations (left) of 30 PYR cell inputs onto an OLM cell model. Simulated voltage traces (right; as in A) but with a scaling up of PYR cell inputs alone instead of I-S3 cell inputs alone. D-F: Blown up versions of the traces highlighted by the stars in A-C, respectively. In all cases, the models are given somatically injected current to generate spiking at 7.25 Hz (cell 1: 49 pA; cell 2: 45 pA) to generate the mean baseline OLM cell spike rates reported by Tyan et al. (15). This figure displays results for cell 1. Results for cell 2 are similar (**OSF Figure 2)**.

##### Phase response curve (PRC) analyses

The control of OLM cells by I-S3 cells has been suggested to be due to PIR mechanisms (15). That is, inhibitory inputs would activate biophysical currents such as *I*_*H*_ in OLM cells which could lead to spiking. Considering this, we are in a position to take advantage of our detailed OLM cell models that were developed in a “neuron-to-model matched” (NMM) fashion (27) regarding its morphology, passive properties, and biophysical currents. These are currently the most biophysically, detailed OLM cell models and using them serves as a biological proxy for exploring PIR mechanisms in a comprehensive fashion. We do this by turning to phase response curve (PRC) analyses, which have been used to study morphologically detailed neuron models in the past (36). If PIR mechanisms are in play, then one would expect to observe that I-S3 cells could cause phase advances and not phase delays of OLM cell spiking.

To compute PRCs as shown in **Figs. 3B,C, 4A**, we compute the phase response (ΔΦ) for perturbations given at different phases of the cycle. We obtain the interspike interval of the two spikes preceding the perturbation (*T* 0) and the interspike interval between the last spike preceding the perturbation and the first spike following the perturbation (*T* 1). We calculate the phase response as follows:

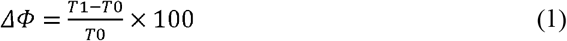

In this sense, the phase response is normalized to 1 and converted to a percent. A negative value means a phase advance, or a shortening of the interspike interval. A positive value means a phase delay, or a lengthening of the interspike interval. We compute the phase responses where the zero perturbation is defined as the time when the spike crosses a threshold of -20 mV during its upstroke.

**Figure 3:**
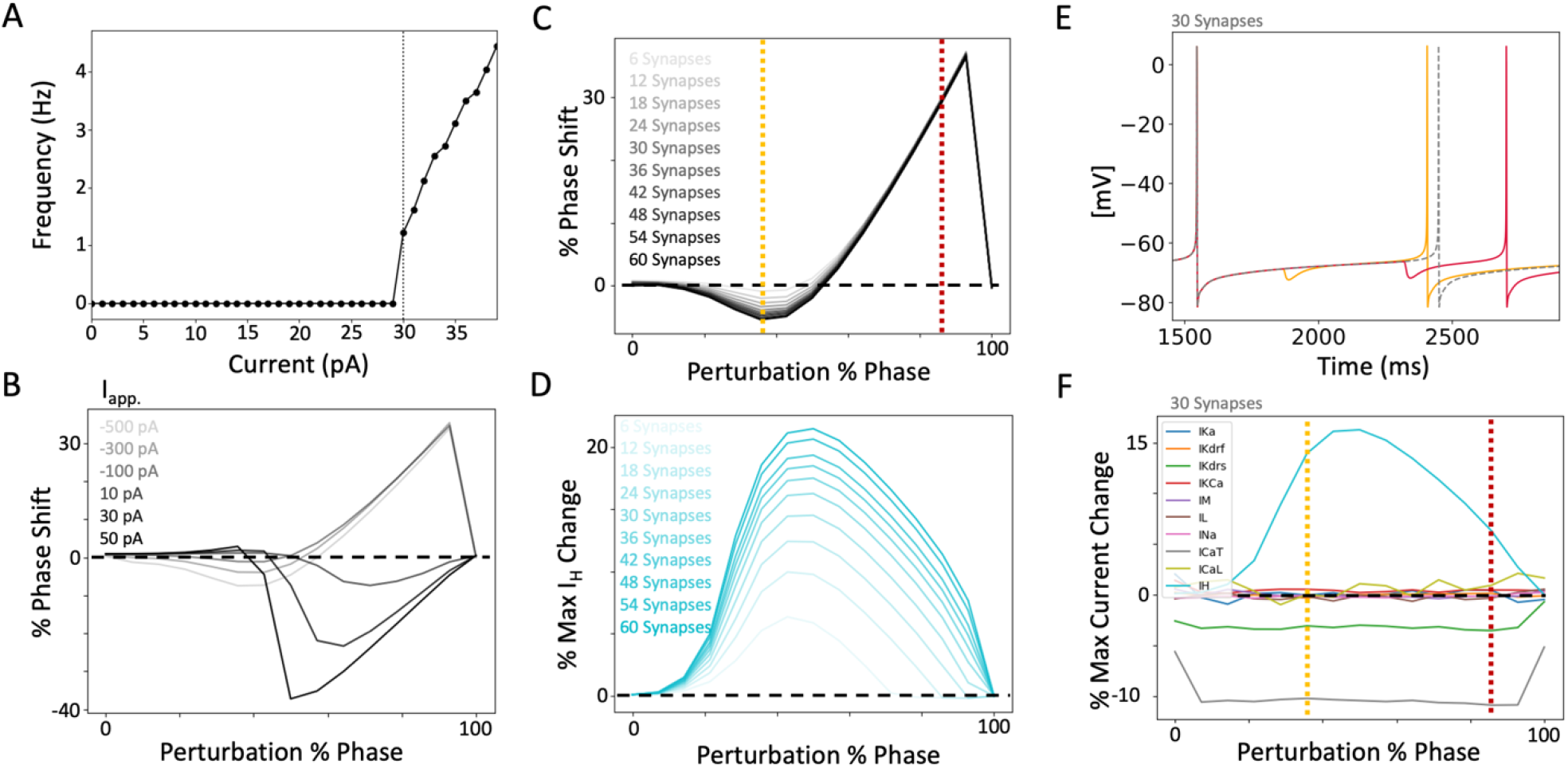
PRC exploration illustration. **A:** Relationship between holding current and resulting spike rates in cell 1. Dashed line indicates the holding current level used for generating baseline spiking that is used to examine phase responses in panels B-F. B: Resulting PRCs due to large negative and small positive 3ms square current pulses (−500, -300, -100, 10, 30, 50 pA) being moved from earlier (0%) to later (100%) phases of an inter-spike interval cycle. More positive phase shifts indicate larger phase delays. C: Resulting PRCs due to inhibitory perturbation (6 to 60 I-S3 cell synapses) being moved from earlier (0%) to later (100%) phases of an inter-spike interval cycle. Example perturbation phases at which an advance and a delay can be seen are indicated by the yellow and red dashed lines, respectively. D: Perturbation phase-dependent changes in peak *I*_*H*_ amplitude for different numbers of I-S3 cell synapses. E: Voltage traces at baseline without perturbation (dashed), with an advance-causing perturbation of 30 I-S3 cell synapses (yellow), and with a delay-causing perturbation of 30 I-S3 cell synapses (red). Colors correspond with the perturbation phases highlighted by the colored dashed lines in C and F. F: Perturbation phase-dependent changes (30 I-S3 cell synapses) in peak current amplitude for each ion channel present in the model. Currents are colored as follows (same order as in legend): *I*_*KA*_ (blue), *I*_*Kdr f*_ (orange), *I*_*Kdrs*_ (green), *I*_*KCa*_ (red), *I*_*M*_ (purple), *I*_*L*_ (brown), *I*_*Na*_ (pink), *I*_*CaT*_ (grey), *I*_*CaL*_ (yellow), and *I*_*H*_ (cyan). The measurements shown here are extracted from current traces obtained from the first dendritic compartment adjacent to the soma since calcium channels are not present in the somatic compartments.

**Figure 4:**
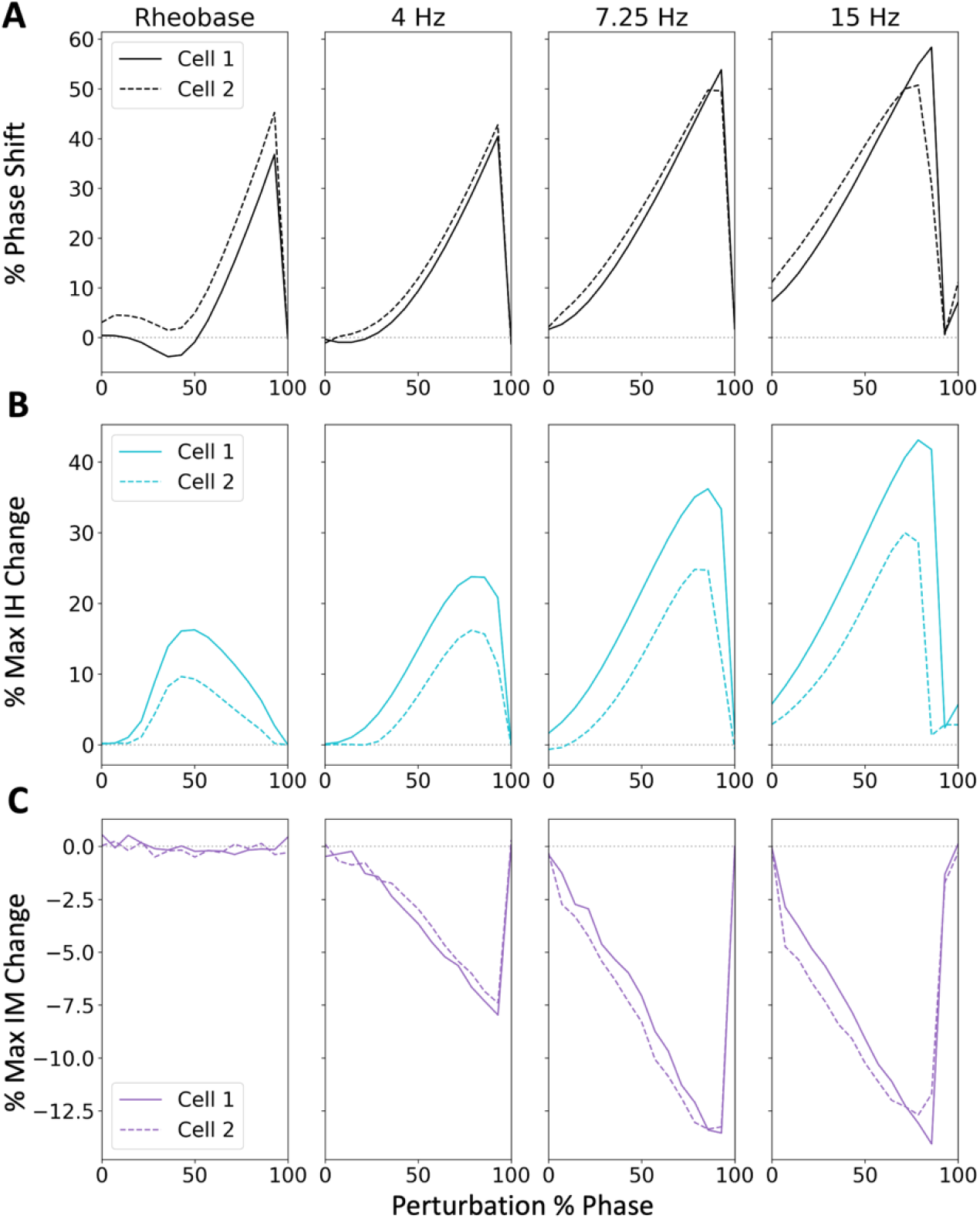
*I*_*H*_ -modulated phase advances only occur when spiking is near rheobase. **A:** PRCs for cell 1 (solid) and cell 2 (dashed), during rheobase, 4 Hz, 7.25 Hz, and 15 Hz OLM cell spiking, using a perturbation of 30 I-S3 cell synapses. B: Perturbation phase-dependent changes in peak *I*_*H*_ amplitude for cell 1 (solid) and cell 2 (dashed). C: Perturbation phase-dependent changes in peak *I*_*M*_ amplitude for cell 1 (solid) and cell 2 (dashed). It is clear that inhibitory perturbations alone mostly just cause phase delays in OLM cell models.

To compute the change in biophysical currents (the same calculation used for each current type; see **Figs. 3D,F, 4B,C**) caused by the perturbation, we obtain the peak current amplitude generated in the period from the 2nd last spike preceding the perturbation to the perturbation time (*I*0), and the peak current amplitude generated in the period from the perturbation time to the 2nd spike following the perturbation (*I*1). Percent change in peak current is calculated as follows:

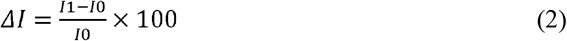

As such, positive values indicate percent increases in peak current amplitude, and negative values indicate percent decreases in peak current amplitude.

#### In vivo model exploration

##### *In vivo* setup

It is known that the large amount of synaptic bombardment in the behaving animal *in vivo* changes the response profile of individual neurons (37). In previous work using I-S3 cell models (32), we had developed techniques for performing high-resolution parameter searches in parallel on the Neuroscience Gateway (NSG) for high-performance computing (38) to find synaptic input parameter combinations (i.e. excitatory and inhibitory numbers of synapses and spike rates) that could generate *in vivo*-like (IVL) states for the given cell model. IVL states were characterized by metrics that ensured, for example, large enough fluctuations of subthreshold membrane potential activities and interspike interval coefficient of variations. In Guet-McCreight and Skinner (33), we used similar techniques to obtain IVL states for OLM cells. We found that IVL states resulted in synaptic parameter values for cell 1 of 1268 excitatory synapses firing at 1.6 Hz and 1254 inhibitory synapses firing at 8.7 Hz, and for cell 2, 1503 excitatory synapses firing at 1.5 Hz and 1532 inhibitory synapses firing at 8 Hz, distributed throughout the dendritic tree.

#### Spiking resonance

##### Setup for *in vitro* spiking resonance

To examine spiking resonance in *in vitro* states, we generate 50 different baseline spike rates (*f*_*B*_’s) in our OLM models by applying a range of holding currents from 30-152.5 pA (cell 1) and 22-144.5 pA (cell 2). Note that plots of the frequency-current relationships of these models have been reported previously (33). This elicits a range of *f*_*B*_’s from about 1-35 Hz in both models. Specifically, for cell 1, we obtain a *f*_*B*_ range of 1.14-35.72 Hz, and a range of 0.95-34.80 Hz for cell 2, for the holding current ranges used. Each of these models is then subjected to a range of inputs at different input frequencies (0.5 -30 Hz) to determine spike resonance frequencies.

##### Setup for *in vivo* spiking resonance

To examine spiking resonance in *in vivo* states, we also generate 50 different *f*_*B*_’s in our OLM models. However, this is done by changing the random seed that controls placement of synapses and presynaptic spike times rather than by changing the holding current since *in vivo* one is not injecting tonic current into the soma. This generates different *f*_*B*_’s (across random seeds), though not with as wide of a range as changing the holding current in the *in vitro* state. We obtain 50 *f*_*B*_’s by using 50 different random seeds. Specifically, for cell 1, we obtain a *f*_*B*_ range of 11.82-15.83 Hz, and a range of 9.34-13.57 Hz for cell 2. Each of these models is then subjected to a range of inputs at different input frequencies (0.5 -30 Hz) to determine spike resonance frequencies.

##### Spiking resonance calculation

For each of the models, we compute the power spectral density (PSD) of the spike train (1’s for spikes and 0’s for no spikes, using a -20mV threshold for spikes) without modulatory inputs (i.e., at *f*_*B*_). The PSD is computed in python using the welch function available as part of the scipy module: `signal.welch(signal, fs=1/dt, scaling=‘density’, nperseg=20000)`. We then apply a series of different input frequencies (*f*_*I*_) using different input populations, and the PSD is computed following each input frequency.

The different input populations used are: 30 frequency-modulated I-S3 cell inputs, or 30 frequency-modulated PYR cell inputs, or 30 frequency-modulated inputs from each input populations (i.e., I-S3, MS, BIS, and PYR cells). For the latter, the start time delays from each other are based on their relative timing during theta rhythms (16, 39, 40). Specifically, these start times (phases) relative to a 125 ms theta cycle are the following: I-S3, 45°; MS, 225°; BIS, 270°; PYR, 270 °. We note that the choice to use 30 I-S3 cell synapses is because at this number of synapses, I-S3 cell synaptic activation can exhibit a sufficient level of synaptic control over OLM cell spiking at 5 Hz in both models (see **Fig. 2)**. The choice to use the same numbers of PYR cell, MS cell, and BIS cell synapses is simply to keep the numbers equivalent in the absence of appropriate experimental data obtained in this context to constrain these numbers.

To gauge spike frequency resonance, we use a ratio to baseline measurement (i.e., ‘baseline ratio’ or δ *PSD*), which is computed by dividing the PSD at the *f*_*I*_ by the PSD at the *f*_*I*_ in the corresponding baseline trace (i.e. without the modulatory inputs):

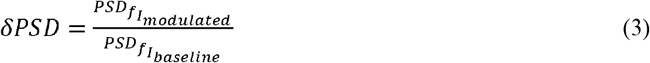

In other words, it is a measurement of how much the PSD at the *f*_*I*_ changes relative to baseline once modulatory inputs are added. A value of 1 would therefore indicate that there is no change in PSD, values less than 1 indicate a decrease in PSD, and values greater than 1 indicate an increase in PSD. The magnitude of this metric therefore indicates the effectiveness of synaptic inputs at entraining spiking to a prescribed input frequency. Spike resonance (*f*_*R*_) is defined as the *f*_*I*_ generating the largest baseline ratio.

#### In vivo theta rhythm exploration setup

We use our models to consider an *in vivo* context as found in (16). We run simulations using the full IVL scenarios + 8 Hz modulatory inputs from all input population (I-S3, MS, BIS, and PYR cells). As done in Luo et al. (16), we add a small amount of noise in all of the theta-timed inputs, as this is both more realistic than having perfectly-timed inputs, and enhances theta recruitment.

## Results

Focusing on I-S3 cell inhibitory control over OLM cells, we perform detailed computational explorations of OLM cell models considering *in vitro* and *in vivo* contexts. These explorations represent virtual network scenarios as the OLM cell is either depolarized with holding current injections (*in vitro* state) or virtually embedded in a network where it is receiving much synaptic bombardments (*in vivo* state). In **Fig. 1** we show an example of output in *in vitro* (A) and *in vivo* (B) contexts along with the biophysical currents that underlie the outputs.

### I-S3 cell control over OLM cell spiking – reproducing in vitro experimental observations

*In vitro*, optogenetic activation of CR+ cells, which includes I-S3 cells, can preferentially control the timing of OLM cell spiking at 5 and 10 Hz, but not 1 and 20 Hz. Tyan et al. (15) considered that this was achieved through PIR spiking mechanisms. However, because other inhibitory synapses and excitatory synapses were not blocked in these experiments, it is unclear whether this is due to network effects or to intrinsic, biophysical OLM cell ion channel properties that promote PIR spiking.

We first note that following negative long-square, high amplitude current injections, OLM cells are capable of exhibiting rebound spikes (27). Our OLM cell models also exhibit this feature (Cell 1 threshold: -220 pA; Cell threshold 2: -230 pA), primarily through activation of *I*_*H*_ and subsequent activation of *I*_*Na*_ (**OSF Figure 1** on https://osf.io/28r4s/). However, it is unclear if synaptic inhibition alone is sufficient to trigger this same phenomenon. As well, it is unknown whether and how findings obtained *in vitro* would translate to *in vivo* contributions in a behaving animal.

#### I-S3 cell input populations alone cannot entrain OLM cell spiking as observed experimentally

There are several ways in which I-S3 cells could manifest their control over OLM cells. If it is primarily through PIR spiking, this implies that inhibitory perturbations from I-S3 cells to OLM cells essentially speed up the onset of an OLM cell spike – a phase advance due to intrinsic OLM cell properties such as *I*_*H*_, *I*_*CaT*_, and *I*_*M*_ that would interact with the incoming inhibitory perturbations. There are other ways in which I-S3 cells could influence OLM cells in the hippocampal circuitry. I-S3 cells could be controlling the timing of OLM cell spiking by suppressing and delaying OLM cell spiking. This would require OLM cells to be spiking faster than the frequency at which I-S3 cells are activated to exert their control. During the *in vitro* experimental recordings, the average spike rate of the OLM cells is 7.3 Hz and OLM cells could be recruited to spike at 10 Hz photo-activation patterns. Thus, simple suppression by I-S3 cells seems unlikely to be the case.

Let us start by examininghowI-S3 cell synaptic inputs alone could control OLM cell output. Using optimized synaptic parameters of I-S3 cell inputs to our detailed OLM cell models, we explore a range of different numbers of synapses when I-S3 cells are activated at different frequencies as was done in Tyan et al. (15). An example of 30 I-S3 cell synapses distributed on one of the OLM cell models is shown in **Fig. 2A**.

We apply I-S3 cell synaptic inputs alone with rhythmic spiking at 1 Hz, 5 Hz, 8 Hz, 10 Hz, and 20 Hz and using 0 to 60 synapses. The resulting output from one of the OLM cell models (cell 1) is shown in **Fig. 2A**. Output for cell 2 is similar and is provided in **OSF Figure 2** on https://osf.io/28r4s/. In looking at the spike traces across both cell 1 and cell 2, we observe that entrainment by the I-S3 cell synapses occurs primarily at 5 Hz when there is a large enough number of frequency-modulated synapses. However, at higher frequencies, I-S3 cell inputs only lead to spiking on every other cycle. For example, at 10 Hz stimulation with 30 I-S3 cell synapses, the OLM cell models spike at 5 Hz. This is in stark contrast with the experimental results that show that OLM cells spike at 10 Hz when receiving 10 Hz inhibition from CR+ cells (15). These modeling results show that I-S3 cell synaptic inputs delay the spiking of OLM cells on each cycle, which is contrary to what one would expect if PIR mechanisms in OLM cells were the main way by which I-S3 cells were exerting their control of OLM cell firing. We also tested the inclusion of inhibitory input from other populations onto OLM cells [e.g., MS cell inputs, which were reported by Chamberland et al. (13) to have comparatively larger IPSC amplitudes than I-S3 cell inputs], as well as clustering I-S3 cell synapses on distal or proximal dendrites [i.e., since I-S3 cell IPSC amplitudes from Chamberland et al. (13) suggest preferred distal localization of I-S3 cell synapses]. These manipulations did not lead to better spike entrainment at 10 Hz frequencies (not shown). Further, these explorations suggest that any other purely inhibitory pathway, e.g., a ‘I-S3 to BIS to OLM cell’ pathway, would also lead to spiking delays, and not be able to reproduce the experimental observations of Tyan et al. (15). We note that IPSP magnitudes (**Fig. 2D**) are comparable to those generated experimentally during wide-field optogenetic stimulation of CR+ cells [i.e., Fig 8C1-C4 in Tyan et al. (15)], likely because the I-S3 cell input conductances in our models were optimized to capture the amplitudes of I-S3 to OLM cell connections reported by Chamberland et al. (13). However, this blown-up trace (**Fig. 2D**) also highlights that the full measured amplitudes of the IPSPs are dependent on their timing relative to the after-spike hyperpolarization portion of the trace, which can occlude the IPSP almost completely.

#### I-S3 cell inputs together with PYR cell inputs can entrain OLM cell spiking as observed experimentally

To reproduce the *in vitro* experiments, it would seem to be the case that I-S3 cells do not simply suppress and delay inhibitory cell spiking without PIR spiking. However, since excitatory synapses were not blocked in the *in vitro* experiments, it is possible that photoactivation of I-S3 cells might have the general effect of disinhibiting PYR cells, which could then further phase lock the spiking of OLM cells through recurrent excitation (i.e., a ‘I-S3 to OLM to PYR to OLM cell’ pathway). There are additional ways in which this could occur. For example, it is possible that inhibition of BIS cells by I-S3 cell activation could contribute towards disinhibiting PYR cells, which could further augment recurrent excitation to OLM cells in an ‘I-S3 to BIS to PYR to OLM cell’ pathway.

Thus, since inhibitory inputs alone are not able to entrain OLM cell spiking in a manner that is consistent with the *in vitro* experiments, we include PYR cell inputs in a disinhibitory manner to determine whether we could achieve a match with experiment. As we are not explicitly modeling PYR cells, we include frequency-modulated PYR cell synaptic inputs with a delay relative to inhibitory inputs. Also, we add only a few PYR cell synapses (i.e., enough to generate spiking) on the OLM cell models to determine whether we could obtain the level of spike entrainment previously seen in experiment (15). This is shown in **Fig. 2B** for cell 1 (cell 2 in **OSF Figure 2** on https://osf.io/28r4s/) where it can be seen that one-to-one entrainment is possible at 5 and 10 Hz as observed experimentally, but now also at 20 Hz, which is unlike what is seen experimentally in Tyan et al. (15).

The inclusion of frequency-modulated inputs from both I-S3 and PYR cells is ‘virtual’, meaning that we are not explicitly modelling I-S3 and PYR cells in a circuit configuration with the OLM cell models. That is, in our virtual network simulations here, we include PYR cells inputs at the various frequencies, assuming that they reliably spike at the given input frequency, whether it be 5 or 20 Hz and so on, as there is no explicit modeling of tri-synaptic connectivity between I-S3 cells, OLM cells, and PYR cells. While this is a clear limitation, the alternative to simulate full microcircuits brings about many other aspects to consider and introduces other limitations and caveats (see Discussion). That being said, a possible interpretation of our simulation results is that, experimentally, photo-activation of CR+ cells at 20 Hz may not lead to disinhibited PYR cell spiking at 20 Hz, as modelled here with our virtual networks -at least not enough to excite OLM cells to spike at these higher frequencies. Additionally, wide-field optogenetic stimulation of CR+ cells in Tyan et al. (15) was often capable of generating 2-3 OLM cell spikes. This effect was not captured when scaling up the number of I-S3 cell inputs alone (**Fig. 2A**) or with a minimal number of PYR cell inputs (**Fig. 2B**). In scaling up the number of PYR cell inputs on its own, however, we find that this effect can be replicated (**Fig. 2C & F**). As such, our simulations suggest that wide-field activation of CR+ cells, that includes I-S3 cells, may disinhibit a large enough number of PYR cells to cause a robust recruitment of OLM cells to spike.

### Insights from PRC analyses with biophysical correlates

PRCs are used in studies of oscillatory systems such as neuronal spiking (41, 42) to indicate the change in the cycle period as a function of the phase at which a perturbation is received, thus predicting its response to rhythmic inputs -that is, its entrainment at various frequencies. Thus, we here use PRCs to uncover whether there are any scenarios under which I-S3 cells could cause phase advances of OLM cell spiking and thus indicative of PIR mechanisms. In the previous section where we focused on replicating *in vitro* experiments (15), we did not find support for PIR mechanisms contributing to I-S3 cell control of OLM cell firing.

#### Inhibitory perturbations rarely elicit PIR spiking in OLM cell models

In **Fig. 3**, we illustrate theoretical PRC explorations. **Fig. 3A** shows the spiking frequency of cell 1 for a given amount of somatically injected tonic current. At 30 pA injected current, as indicated by the dashed line, the cell fires at about 1 Hz. The PRCs generated for the cell firing at this frequency are given in **Fig. 3B & 3C**, in response to different positive and negative 3ms square current pulses at the soma, and different numbers of I-S3 cell synapses spread throughout the dendrites, respectively. Both negative current pulses with magnitudes up to -500 pA (**Fig. 3B**) and up to 60 inhibitory synaptic inputs (**Fig. 3C**) could generate phase advances at early phases of the interspike interval. In comparison, much smaller amplitude positive current pulses caused large phase advances at later phases of the interspike interval (**Fig. 3B**), and similar PRC shapes could be obtained with PYR cell synapses, but with conductance values that are much smaller than optimized PYR synapse values (not shown). Increasing the inhibitory perturbation past 30 synapses had little effect on further enhancing phase advances (**Fig. 3C**) but was accompanied with a continuous increase in *I*_*H*_ response (**Fig. 3D**). The dashed lines in **Fig. 3C** show that cell 1 can either show a phase advance (yellow) or a phase delay (red) of the subsequent spike if the cell is perturbed at the given phases indicated along the x-axis.

In **Fig. 3E we** show an overlay of the model voltage traces without perturbation (dashed grey), when there is a phase advance (yellow) or a phase delay (red) using a perturbation of 30 inhibitory synapses. As can be seen in **Fig. 3C**, perturbations for this scenario (OLM cell spiking around 1 Hz, close to its rheobase) mostly lead to phase delays, and any possible phase advances are small. **Fig. 3F** shows the changes in all of the biophysical currents in cell 1 due to the phase-dependent perturbations from 30 inhibitory synapses. It is clear that *I*_*H*_ is the main contributor when there is a phase advance given its change relative to all of the other biophysical currents. This change is as an increase which makes sense for *I*_*H*_ contributing to producing a phase advance since it is an inward current. *I*_*CaT*_, and *I*_*Kdrs*_ also are strong contributors here, but showing decreases for either phase advance or delay scenarios.

We compute the PRCs at four different baseline spike rates (rheobase, 4 Hz, 7.25 Hz, 15 Hz) in the OLM cell models (see **Fig. 4A**) using a perturbation of 30 inhibitory synapses. From this, it is clear that phase delays are almost always present in response to inhibitory perturbations, with the smallest phase delay occurring when the perturbation is given during the spike refractory period, and the largest phase delay occurring when the perturbation is just before spike threshold -this aspect is visualized in the voltage traces shown in **Fig. 3E**. Only at the lowest baseline spike rate (rheobase) can phase advances (negative values) occur when the perturbation is near 40% of the inter-spike interval, and these small phase advancements only occur with cell 1, and not with cell 2. This exposes the biophysical heterogeneity in these two OLM cell models, each of which were developed using an NMM approach (see Methods). In particular, given that *I*_*H*_ constitutes the main current increase with the perturbation, and we know that the h-channel conductance is larger in cell 1 relative to cell 2, we can say that this is what underlies the small phase advancement that occurs in cell 1. **Fig. 4B** shows the *I*_*H*_ changes for the four different baseline spike rates and one can see the larger *I*_*H*_ for cell 1 as compared to cell 2. It is important to note that the various channel conductances in the cell 1 and cell 2 OLM cell models were optimized to the specific electrophysiological data, and not arbitrarily set to different values. However, to verify that variations in *I*_*H*_ in cell 1 or 2 models do not affect these results, we adjust the h-channel conductance by ±25% and find little change in the PRCs (**OSF Figure 3** on https://osf.io/28r4s/). Unlike *I*_*H*_, most currents exhibited perturbation phase invariance at rheobase (**Fig. 3F**), and stronger current changes near spike threshold (i.e., 100% phase) when the cells were spiking at higher frequencies, such as with *I*_*M*_ for cell 1 and cell 2 (**Fig. 4C** and see **OSF Figure 4** on https://osf.io/28r4s/ for all current changes).

In examining all of perturbation phase-dependent amplitude shifts of the different currents due to the various ion channel mechanisms (shown in **OSF Figure 4** on https://osf.io/28r4s/), we observe the following: We obtain phase-dependent increases in *I*_*H*_ (cyan), and *I*_*CaL*_ (yellow), and decreases in *I*_*M*_ (purple), which should all enhance the likelihood of PIR spiking (i.e. more inward current and less outward current). We note that maximal *I*_*CaT*_ (grey) is decreased following the inhibitory perturbation, regardless of the perturbation phase, but is steadily decreased less at later phases of the interspike interval for base firing rates of 4 and 7.25 Hz. A phase-dependent increase, albeit a small one, in 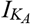 (blue), however, would counteract rebound firing since 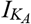 is known to suppress PIR spiking (29). It should be noted that contributions from 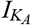 are large in these models, which could partially account for the lack of inhibitory perturbation-dependent phase advances. Other currents (*I*_*Kdr f*_ : orange, *I*_*Kdrs*_: green, *I*_*KCa*_: red, *I*_*Na*_: pink, *I*_*L*_: brown) do not show appreciable inhibitory perturbation phase-dependent changes in maximal magnitude. Comparing relative changes across the baseline firing rates and perturbation-dependent changes, enhancements in contributions from 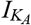, *I*_*Kdr f*_, *I*_*Kdrs*_ and *I*_*KCa*_ all contribute towards the prevention of phase advances from occurring more broadly across higher spike rates. Since 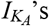 phase-dependence near rheobase is flat compared to its phase-dependence at higher spike rates, this may also account for why some phase advances are permitted near rheobase. Though there is a difference in PRCs across the two models, mainly at the rheobase baseline spike rate, the ion channel current responses are consistent for both OLM cell models. Altogether, these results suggest that inhibitory inputs alone, via I-S3 cells, are unlikely to be sufficient to elicit PIR mechanisms in OLM cells. Rather, the modeling predicts that I-S3 cell activation suppresses OLM cell spiking (i.e., causes phase delays).

### I-S3 cell inhibitory inputs and spike resonance in OLM cells

#### *In vitro* spike resonance at theta frequencies occurs for excitatory but not inhibitory inputs

From our PRC analyses and simulations of the *in vitro* experiments of Tyan et al. (15), it seems clear that I-S3 cells likely do not control OLM cell firing through PIR mechanisms since small phase advances were obtained in a very limited fashion (rheobase for cell 1 only). We have so far limited our explorations to a few input frequencies and baseline spike rates. From these results it is clear that inhibitory modulation depends on the baseline firing rate of the OLM cell (*f*_*B*_), as well as the frequency of the incoming frequency-modulated synaptic inputs (*f*_*I*_), though the precise relationship between the two is unclear. Let us now explore a wide range of baseline firing rates in OLM cells and apply a wide range of frequency-modulated inputs to determine the scenarios under which preferred firing (i.e., spike resonance) could occur, and whether it includes theta (3-12 Hz) frequencies. We do this by incrementally changing the holding current to elicit different *f*_*B*_, while at the same time submitting the model at each holding current to a barrage of different input frequencies. We note that this range of baseline spike rates includes the range of spike rates that have been reported for OLM cells *in vivo* across different behavioral and network states (43, 44). In this way we can establish a ‘ground-truth’ regarding preferred output responses of OLM cells in the face of various input frequencies, i.e., an OLM cell spiking resonance, given different baseline spike rates in an *in vitro* context.

As noted earlier, the *in vitro* context simply means that the OLM cells are not bombarded with excitatory and inhibitory synaptic inputs. In this context, we apply the various holding currents to the somatic compartment (see **Fig. 1A**), and rhythmic synaptic input activations are applied to dendritic compartments. In **Fig 5**, we show the spike resonant frequencies (*f*_*R*_) of cell 1 when receiving inhibitory inputs from I-S3 cells (**Fig. 5A-D**) or excitatory inputs from PYR cells (**Fig. 5E-H**). Results from inputs across all populations (i.e., including MS and BIS inputs) are shown in **OSF figure 5** on https://osf.io/28r4s/, as well as for both cell 1 and cell 2. When receiving only I-S3 cell inhibitory inputs (**Fig. 5A**), *f*_*R*_ is largely dependent on *f*_*B*_ (**Fig. 5B**; cell 1 *f*_*R*_: 15.37 ± 8.76 Hz; cell 2 *f*_*R*_: 15.23 ± 8.88 Hz). That is, if the OLM cell is firing at a higher rate, then *f*_*R*_ is also higher. This can be seen more clearly in **Fig. 5C**, which also highlights that *f*_*R*_ can also occasionally be larger than *f*_*B*_, despite having spike rates at *f*_*R*_ that are always suppressed compared to *f*_*B*_ (**Fig. 5D**; i.e., consistent with the phase delays due to inhibitory perturbations reported in previous sections). One explanation for this is that when the baseline ratio is enhanced in those cases, the spikes that are being suppressed are spikes that occasionally fall in phase with the modulatory inputs. In this sense, the modulatory inhibitory inputs are keeping the spiking entrained through suppression. This can happen regardless of whether *f*_*R*_ is greater than or lesser than *f*_*B*_, and depends on how the intrinsic spike train aligns with the modulatory input spike trains.

**Figure 5:**
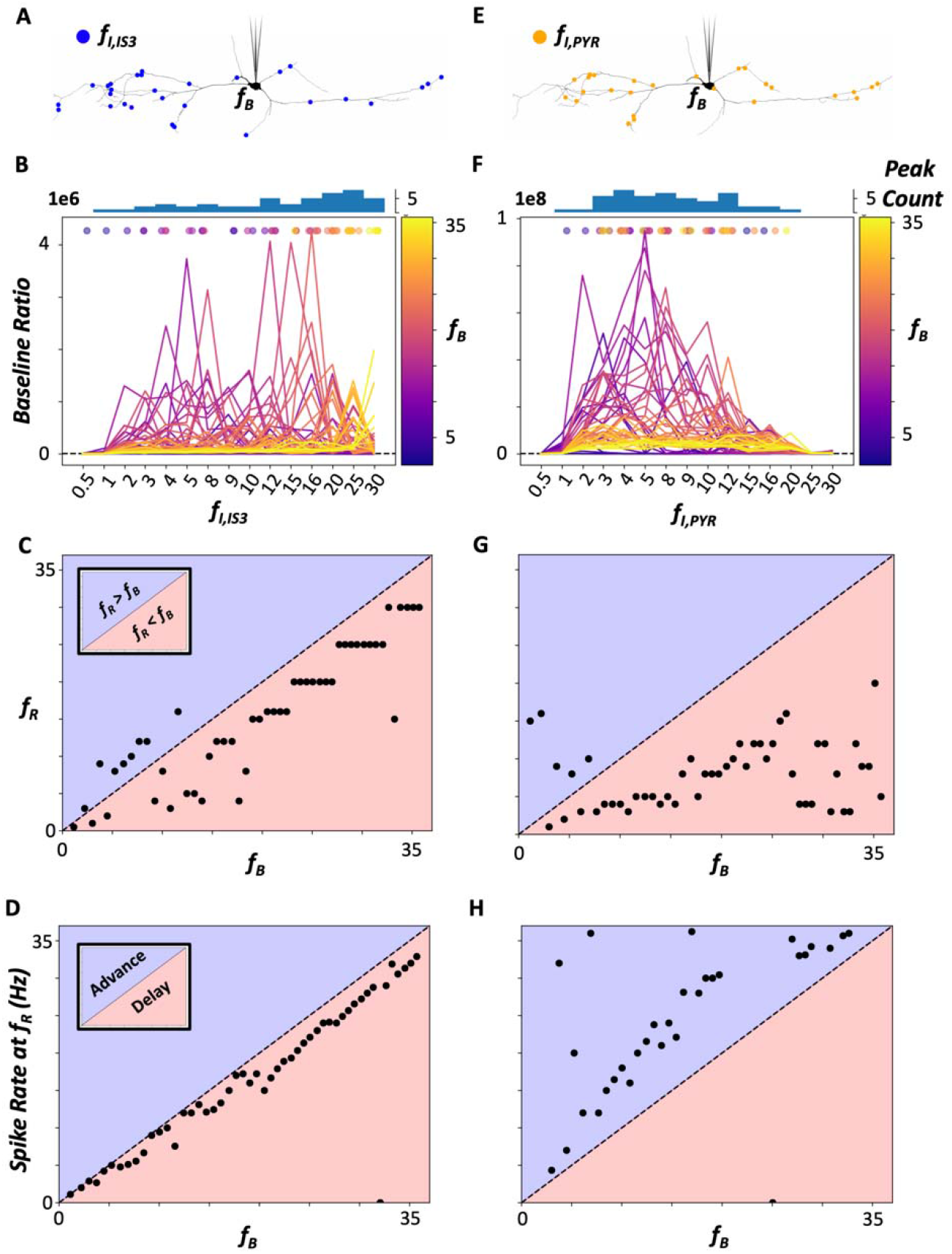
Cell 1 *in vitro* spike resonant frequencies due to inhibitory inputs, but not excitatory inputs, are dependent on the baseline spike rate. A: Example synaptic locations of 30 I-S3 cell modulatory inputs during somatic current injections to generate different *f*_*B*_. B: Baseline ratios computed across different I-S3 cell modulation frequencies, *f*_*I*_, and different holding currents (as shown by the range of different baseline frequencies, *f*_*B*_, indicated by the colorbar). The colored dots and histograms plotted above the traces indicate the peak/resonant frequencies and distributions across *f*_*B*_ values, respectively. The dashed black line indicates a baseline ratio of 1 (note that the y-axis scale is large, so this line appears very close to 0). Values larger/smaller than 1 indicate an increase/decrease in the PSD at *f*_*I*_ . C: Resonant frequencies, *f*_*R*_, plotted against *f*_*B*_. Dots in blue/red areas indicate simulations where *f*_*R*_ is greater/lesser than *f*_*B*_. D: Spike rate at the *f*_*R*_ plotted against *f*_*B*_. Dots in blue/red areas indicate simulations where the *f*_*R*_ spike rate is increased/decreased compared to *f*_*B*_ (i.e. consistent with phase advances/delays). E-H: Same as in A-D, but with 30 PYR cell modulatory inputs instead.

Overall, these results make sense given the findings from the previous section showing that I-S3 cell synapses alone rarely lead to phase advances. Specifically, recruitment of OLM cell spiking to certain inhibitory input frequencies is largely baseline spike rate-dependent because inhibitory perturbations mostly just cause phase delays. For example, if the baseline spike rate is much smaller than the input frequency, inhibitory perturbations will only cause suppression of spikes, and the PSD at the input frequency is more likely to drop. Likewise, if the baseline spike rate is much larger than the input frequency, then inhibitory perturbations will have minimal effects on spiking, as they may only suppress a small fraction of spikes. Thus, the spike resonant frequency is largest when the baseline spike frequency is near the inhibitory input frequency. These tests were also performed using MS inputs alone (not shown), which have larger IPSC amplitudes, and similar results are obtained.

In contrast, OLM cell spike resonance due to excitatory PYR cell inputs (**Fig. 5E**) is consistently in the 2-15 Hz input frequency range, regardless of baseline spike frequency (**Fig. 5F-G**; cell 1 *f*_*R*_: 7.8 ± 4.28 Hz; cell 2 *f*_*R*_: 4.38 ± 2.48 Hz). However, the magnitude of the baseline ratio at *f*_*R*_ is dependent on baseline spike frequency, with larger baseline spike frequencies having smaller baseline ratios at the *f*_*R*_ (**Fig. 5F**). In all cases the presence of excitatory inputs causes increases in spike rates, which is consistent with phase advances (**Fig. 5H**). Similar findings are seen when including all input populations (see **OSF Figure 5** on https://osf.io/28r4s/), suggesting that any theta frequency (3-12 Hz) spike resonance *in vitro* is largely dictated by excitatory inputs when they are present and not by inhibitory inputs alone. We note that using a lower number of PYR cell synapses could affect these results, but not likely in a drastic way given that OLM cell entrainment is strong even when using a small number of PYR cell synapses (see **Fig. 2C**).

#### *In vivo*, theta frequency spike resonance can occur with inhibitory but not excitatory inputs

We have previously simulated synaptic bombardment conditions on OLM cells to generate *in vivo*-like (IVL) states. This includes a random spread of synaptic locations from different input populations leading to irregular firing -an illustration is shown in **Fig. 1B**. We now investigate I-S3 cell control over OLM cells during IVL states. Since we have already investigated I-S3 cell control over OLM cells in an *in vitro* context, we are able to compare whether or not the same level of control over OLM cell spiking might apply *in vivo*. We use different random seeds to obtain a range of *f*_*B*_’s, and we note that the range of frequencies is not as expansive as *in vitro* as the level of control is limited (i.e., we are not simply injecting tonic somatic currents).

We perform the same spike resonance tests as in **Fig. 5**, but now under this IVL state (Cell 1: **Fig. 6A,E**). We see across all input population types (I-S3 cells and PYR cells) that there is a rightward shift in the resonant frequency distributions, relative to *in vitro*, towards higher values where more if not all *f*_*R*_ values are larger than the *f*_*B*_ values (**Fig. 6B,F**; cell 1 *f*_*R, I*_−_*S*3_: 15.92 ± 5.84 Hz; cell 2 *f*_*R, I*_−_*S*3_: 18.07 ± 7.74 Hz; cell 1 *f*_*R, PYR*_: 27.42 ± 3.43 Hz; cell 2 *f*_*R, PYR*_: 25.6 ± 5.31 Hz). That is, even when considering the smaller range of the baseline spike rates, the resonant frequencies are shifted towards higher input frequency ranges (**Fig. 6C,G**). We note that, as in the *in vitro* case, there is no change in the suppressing effects of inhibitory inputs (i.e., PIR spiking remains absent) or the excitable effects of PYR cell inputs on spike rates (**Fig. 6D,H**). The shift towards higher *f*_*R*_ values than in the *in vitro* case (i.e., **Fig. 5C,G** vs. **Fig. 6C,G)** is possibly because of different aspects associated with the IVL state, including irregular spiking patterns (i.e., spiking is no longer periodic; **Fig. 1B**), as well as a reduction in sensitivity due to a decrease in input resistance. In other words, irregular spike patterns at baseline can allow larger variabilities in *f*_*R*_ values depending on how spike times align with incoming modulatory excitatory and inhibitory inputs. As well, decreased input resistance can increase the required magnitude of excitatory and inhibitory currents needed to modulate the model cell firings. This can be brought about by higher input frequency modulation, thus explaining the increased *f*_*R*_ values. We note again that cell 2 exhibited similar *f*_*R*_ profiles to cell 1, and similar to the *in vitro* case, inclusion of all input populations as modulatory inputs exhibited similar *f*_*R*_ profiles to having just PYR cell inputs (**OSF Figure 6** on https://osf.io/28r4s/).

**Figure 6:**
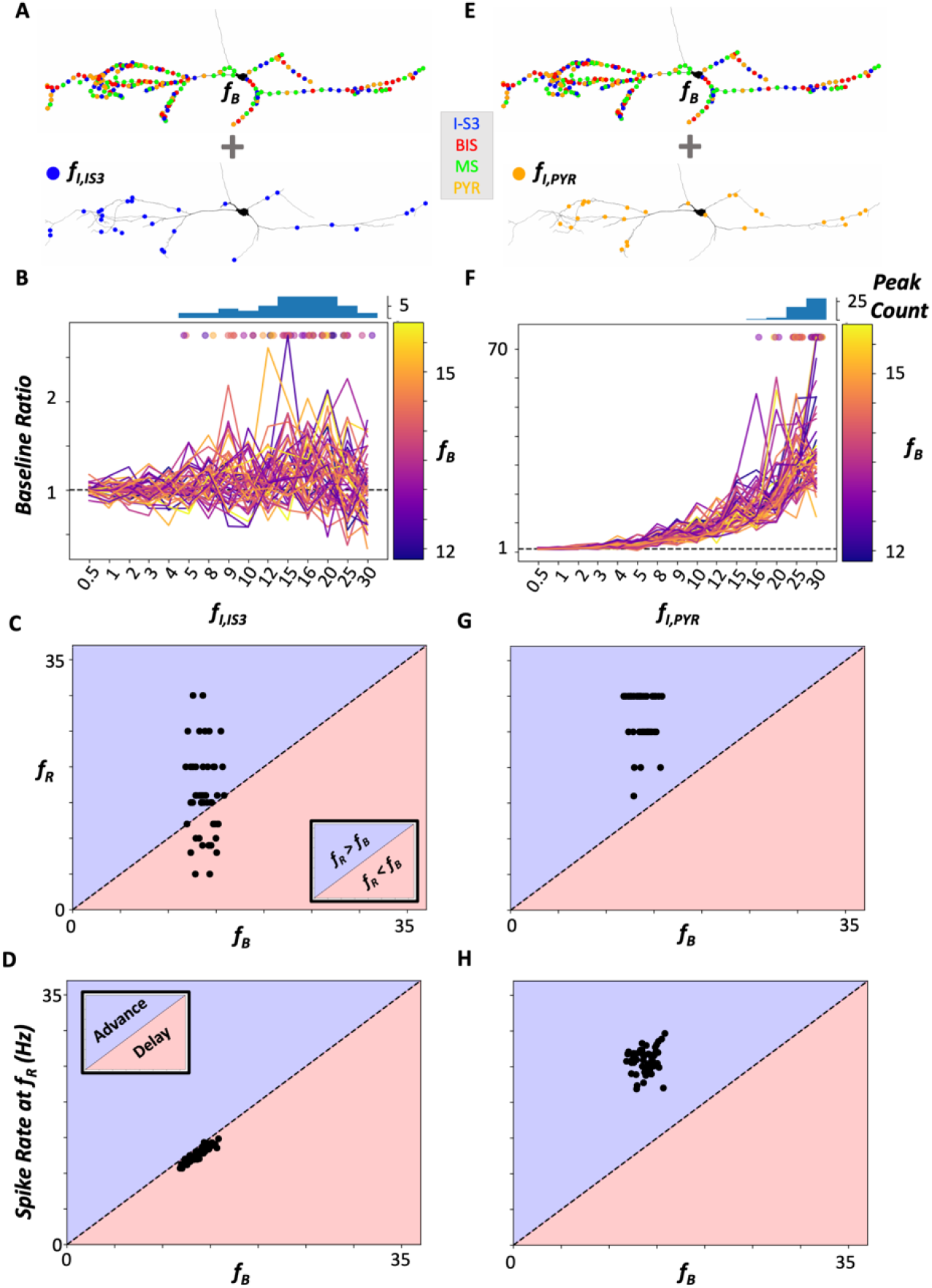
Cell 1 *in vivo* spike resonant frequencies are in theta frequency ranges only for inhibitory inputs. A: Example synaptic locations of IVL synapses (top) and 30 I-S3 cell modulatory inputs (bottom). B: Baseline ratios computed across different I-S3 cell modulation frequencies, *f*_*I*_, and different randomized IVL states (as shown by the *f*_*B*_ range, indicated by the colorbar). The colored dots and histograms plotted above the traces indicate the peak/resonant frequencies and distributions across *f*_*B*_ values, respectively. The dashed black line indicates a baseline ratio of 1. Values larger/smaller than 1 indicate an increase/decrease in the PSD at *f*_*I*_ . C: Resonant frequencies, *f*_*R*_, plotted against *f*_*B*_. Dots in blue/red areas indicate simulations where *f*_*R*_ is greater/lesser than *f*_*B*_. D: Spike rate at the *f*_*R*_ plotted against *f*_*B*_. Dots in blue/red areas indicate simulations where the *f*_*R*_ spike rate is increased/decreased compared to *f*_*B*_ (i.e. consistent with phase advances/delays). Note that axis scales are chosen to match the scales in **Fig. 5C-D & G-H** for comparison purposes. E-H: Same as in A-D, but with 30 PYR cell modulatory inputs instead.

When considering the amplitudes of the baseline ratios, the IVL magnitudes (**Fig. 6B,F**) are considerably smaller than the *in vitro* baseline ratios (**Fig. 5B,F**). This observation could be due to an increase in currents in the model once it is put into an IVL state, which generates a lower input resistance and decreased sensitivity to rhythmically-timed inputs. In other words, higher input frequencies will be necessary to elicit spike resonance since there is a decrease in sensitivity. We note that we have previously established that the addition of synaptic bombardment leads to decreases in sensitivity to additional inputs (32), so it is not surprising that spiking resonance in OLM cells will be different during IVL states relative to *in vitro* states.

The results between the two OLM cell models are qualitatively similar, despite differences in morphologies and intrinsic parameters. One potential reason for this is the presence of dendritic *I*_*H*_, and dendritic Na+ and K+ channels in both models, which would promote similar integration of synaptic inputs and action potential propagation and thus generate qualitatively similar results across both models. As well, our simulation results are quite similar between using I-S3 cell inputs or MS inputs (not shown), the latter of which have larger IPSC amplitudes (13). Indeed, there would not be a strict differentiation of incoming inhibitory inputs as modelled in these virtual networks.

### Simulating I-S3 cell inputs to OLM cells in theta-run epochs during behaviour

In previous work, we found that I-S3 cells are activated with a delay relative to theta-run epochs and spike during the rising to peak phases of theta rhythms (16). As well, it was predicted from modelling work that the timing of this phasic preference would be modulated by inputs from entorhinal cortex (EC; rising phase) and CA3 (peak phase). Thus, we investigate the effects of a ramp-up of I-S3 cell activation (i.e., simulating a delay in I-S3 cell activation) on a per theta cycle basis during an ongoing theta rhythm (**Fig. 7**, which shows results for cell 1). We run simulations using the full IVL scenarios + 8 Hz modulatory inputs from all input population (I-S3, MS, BIS, and PYR cells). As schematized in **Fig. 7A**, starting at 2 s into the simulations until the end of the simulation (10 s), we add 7 I-S3 cell input spikes per cycle, which essentially ramps up the inhibition from I-S3 cells gradually. To consider possible shifts in balances of inputs from CA3 and EC, which can present a possible dis-inhibitory circuitry for switching between sensory inputs and memory consolidation inputs, we also explore shifts in the timing of I-S3 cell inputs alone (**Fig. 7A**) or together with a shift in the timing of PYR cell inputs (**Fig. 7E**). Note that synaptic location sites are chosen randomly for each addition, as described previously (32). The rationale behind these simulations is that the majority of I-S3 cell activation occurs near the ends of theta-run epochs, with a delay relative to the timing of activation of other neuron types, and so we want to see the effect of a ramp-up of I-S3 cell inputs to OLM cell spiking during a behavioural context with ongoing theta rhythms. Here we use five different random seeds for IVL synapses, where IVL synapse locations and spike times are re-randomized (e.g., in **Fig. 1B**).

**Figure 7:**
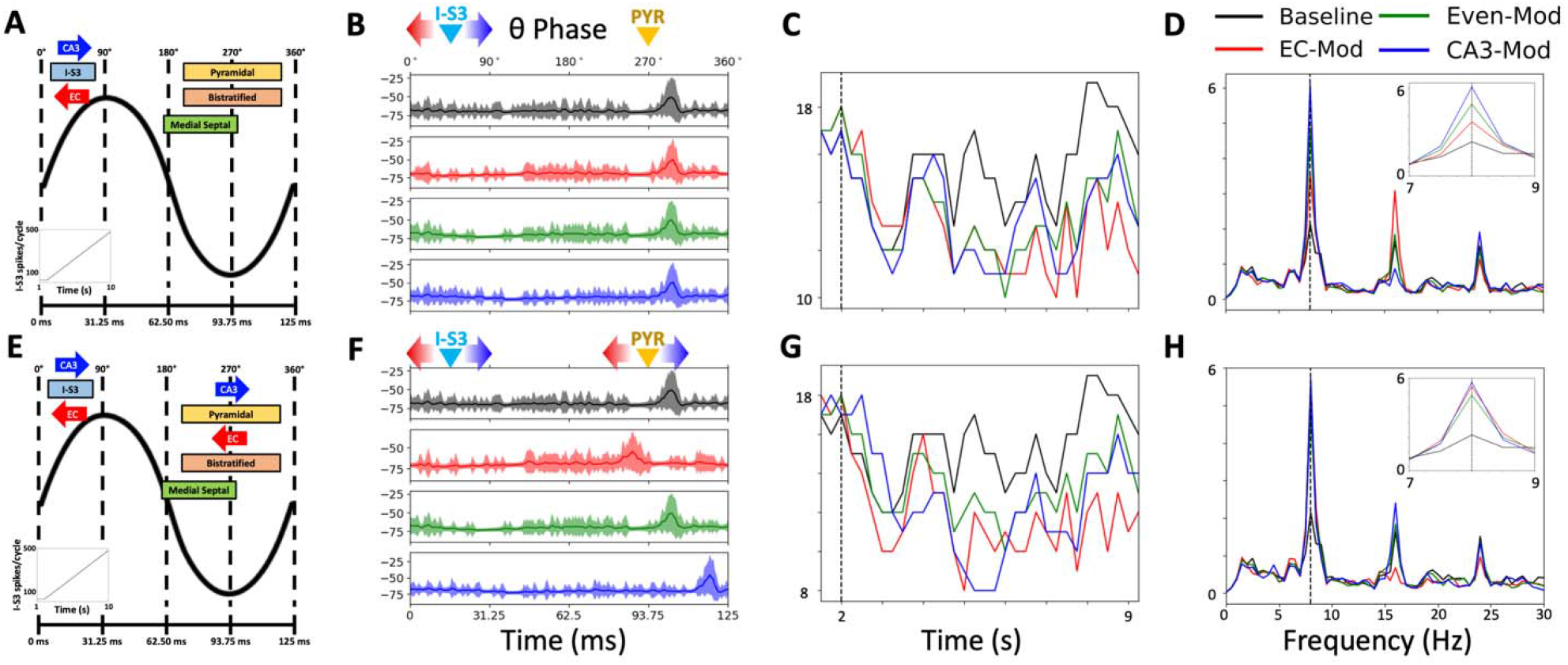
Ramping up I-S3 cell inputs can sharpen and modulate OLM cell recruitment (cell 1). **A:** Schematic of simulations where, on top of IVL inputs, we add theta-timed inputs, a ramp-up of I-S3 cell inputs as each simulation progresses (depicted in the bottom-left subplots) as well as either CA3 (+15.625°) or EC (−15.625°) -modulated shifts in the timing of different input populations. In this first case we impose a shift in the timing of I-S3 cell inputs alone as I-S3 cell inputs are ramped up. B: Plots of the simulated OLM cell model voltage traces averaged across theta cycles. From top to bottom, these plots show the model simulations without a ramp-up (black), with an I-S3 cell ramp-up near the rising phase (red; i.e. stronger CA3 inputs), with an I-S3 cell ramp-up between the rising and peak phases (green; i.e. even inputs from CA3 and EC), as well as with an I-S3 cell ramp-up near the peak phase (blue; i.e. stronger EC inputs), while it is being modulated by 8 Hz frequency inputs from all input population types. To generate these average traces, we take the voltage traces (1,000 to 10,000 ms), split them each into their 72 theta cycles (i.e. 9000 ms/125 ms = 72 cycles), and then compute the average 125 ms theta cycle traces. Shaded areas show the amount of standard deviation above or below the mean. C: Resulting suppression on the spike rate of the OLM cell model throughout the simulations. D: The PSD of the voltage traces before and after applying ramp-ups of I-S3 cell inputs. The inset plot shows the magnified PSD between 7 to 9 Hz. E: In the second case we impose shifts in the timing of both I-S3 cell inputs as well as PYR cell inputs as I-S3 cell inputs are ramped up. F-H: Same as in B-D but with shifts in both I-S3 and PYR cell inputs, as described in E.

#### Ramp-up of I-S3 cell inputs during theta sharpens and modulates OLM cell recruitment

In the simulations where we shift the timing of I-S3 cell inputs alone, the theta-timing of the OLM cell models do not change across any of the conditions (i.e., no ramp-up -black, stronger EC inputs -red, even EC/CA3 inputs -green, or stronger CA3 inputs -blue; **Fig. 7B**). We do see a moderate decrease in the spike rate (**Fig. 7C**), and an increase in the 8 Hz power in the PSD (**Fig. 7D**) across all conditions compared to baseline. More specifically, the enhancement in the PSD is strongest when the timing of the I-S3 cell inputs is shifted towards the peak, corresponding with stronger inputs from CA3 (**Fig. 7D, inset**). This appears to be because inhibition occurring following the peak of theta is the most out-of-phase with the trough of theta, which is when the OLM cell models spike, due to excitation from local PYR cells inputs (**Fig. 7B**). Moreover, this also corresponds to the time at which the I-S3 cell inputs will, on average, be most out-of-phase with the OLM cell spike refractory period, which can allow a stronger *I*_*H*_ -mediated response to inhibition (**Fig. 3**). As such, I-S3 cell inputs alone can sharpen OLM cell recruitment at theta frequencies by suppressing spikes that are out-of-phase with the theta-timing of OLM cell spiking. We note that when tested across five different random seeds, these same results are generated consistently (i.e., the effect of ramped up I-S3 cell inputs causing spike suppression, and the strongest 8 Hz power when modulated by CA3).

In the simulations where we shift the timing of I-S3 cell inputs together with PYR cell inputs (**Fig. 7E**), the theta-timing of the OLM cell models are shifted by the same phasic amount (**Fig. 7F**). The interpretation in these simulations is that a shift in the timing of I-S3 cells due to inputs from either EC or CA3 will shift the dis-inhibitory window for PYR cells, and as such, their phasic timing will shift by the same amount. In these simulations we show that this network effect would also shift the timing of OLM cell spiking. Moreover, since the out-of-phase timing is relative to the phasic timing of when the OLM cell models are spiking, we see a similar PSD magnitude at 8 Hz regardless of the direction in which I-S3 and PYR cell inputs are shifted (**Fig. 7H**). As well, the spike rates are again moderately decreased when compared to baseline (**Fig. 7G**). Again, when tested across five different random seeds, the same results were generated consistently, and similar results were obtained when testing cell 2 (**OSF Figure 7** on https://osf.io/28r4s/).

## Discussion

Mapping identified inhibitory cell types to function and behaviour is a challenging endeavour. To address this, we took advantage of our previous studies interfacing experiment with modeling work of inhibitory cell types in the hippocampus. We focused on how specialized interneuron-selective cells, I-S3 cells, might exert their influence over OLM cells, which have a demonstrated gating control between incoming ‘sensory’ and ongoing ‘memory’ information flow. We used two OLM cell models with parameters fit according to experimental data obtained from the same cell, populations of synapses with parameters specific to cell types that synapse onto OLM cells, PRC theoretical analyses, knowledge of I-S3 cell firing *in vivo*, as well as a methodology for generating *in vivo*-like states.

### How do I-S3 cells control OLM cell spiking in vivo?

From the *in vitro* experimental observation of Tyan et al. (15) in which it was suggested that PIR mechanisms contributed to control of OLM cell spiking by I-S3 cell inhibitory inputs, we considered other possibilities. In simulated *in vitro* scenarios, we found that I-S3 cell-mediated disinhibition of pyramidal cells seemed most likely to be the way in which I-S3 cells exert their influence on OLM cell spike timing. I-S3 cell-mediated suppression of OLM cells would be part of enabling this contribution, but *not* PIR mechanisms, and BIS cells could also contribute in these interactions. Thus, based on our explorations, we predict that the influence that I-S3 cells would have on OLM cells would be through spike suppression followed by enhanced excitation of OLM cells due to PYR cell disinhibition.

Interestingly, when simulating *in vivo* states in OLM cells, only inhibitory inputs, including those from I-S3 cells, could lead to a spike resonance at theta frequencies. This was not the case during simulation of *in vitro* states though -there it was excitatory and not inhibitory inputs that could lead to theta frequency spike resonances. In previous modeling work, it was found that OLM cell spike resonance with inhibitory inputs in simplified, simulated *in vivo* states could occur at high or low theta frequencies depending on whether dendritic h-channels were present or not, respectively (31). Here, we used our NMM OLM cell models that were shown to necessarily have dendritic h-channels (27), a different metric for gauging spike resonance (i.e., the maximal baseline ratio), a series of different baseline spike rates [i.e., instead of just 2.5 Hz, as in Sekulić and Skinner (31)], and synaptic parameters estimated from actual input populations measured experimentally. Based on the results here, we predict that theta frequency spike resonance could occur in OLM cells *in vivo* with incoming inhibitory I-S3 cell inputs, as well as from other rhythmic inhibitory inputs like MS and BIS cell inputs.

From previous work, we know that I-S3 cells exhibit a delay relative to theta-run epochs, with a phasic preference towards the rising/peak phases of theta (16). I-S3 cell activation patterns during running and immobility are also highly variable and even PYR cell-like when compared to other CA1 interneuron types (45), possibly due to mixed modulatory excitation from both CA3 and EC (16), which have delayed activation patterns relative to one another (46) . Simulating a ramp-up of I-S3 cell spiking in our virtual networks, we found that we could obtain a sharpening of the timing of OLM cells during theta rhythms, which could contribute to shaping pyramidal cell place fields (47). In fact, silencing SOM+ cells increases PYR cell burst spiking during place field traversals in awake mice (47), and also un-gates synaptic activation along pyramidal cell apical dendrites (48). In line with this, optogenetic silencing of VIP+ cells, which promotes activation of SOM+ cells, dampens both the reshaping of PYR cell place fields and learning of reward site locations (49). Coincidentally, I-S3 cell activation *in vivo* is significantly less correlated with BIS cell activation than it is with basket cell activation, and even less correlated with OLM cell activation, indicating a suppression of BIS cells and an even heavier suppression of OLM cells when I-S3 cells are activated (45). Previous modeling has also shown that OLM cells, via network pathways that include BIS cells, were key to LFP signal robustness of ongoing intrinsic theta rhythms (50). Thus, I-S3 cell contributions, via OLM and BIS cells, could be essential for the existence of robust theta rhythms.

Based on all of this, our proposition for I-S3 cell contributions is illustrated in **Fig. 8** in a series of steps. We predict that I-S3 cells cause phase delays in OLM cell spiking by spike suppression, as opposed to PIR-mediated phase advances (step 1). This would lead to PYR cell spiking due to disinhibition at a particular phase of theta rhythms (step 2), which would subsequently dictate the timing of the OLM cell spiking through excitation (step 3), leading to re-gating of inputs at distal dendrites (step 4).

**Figure 8:**
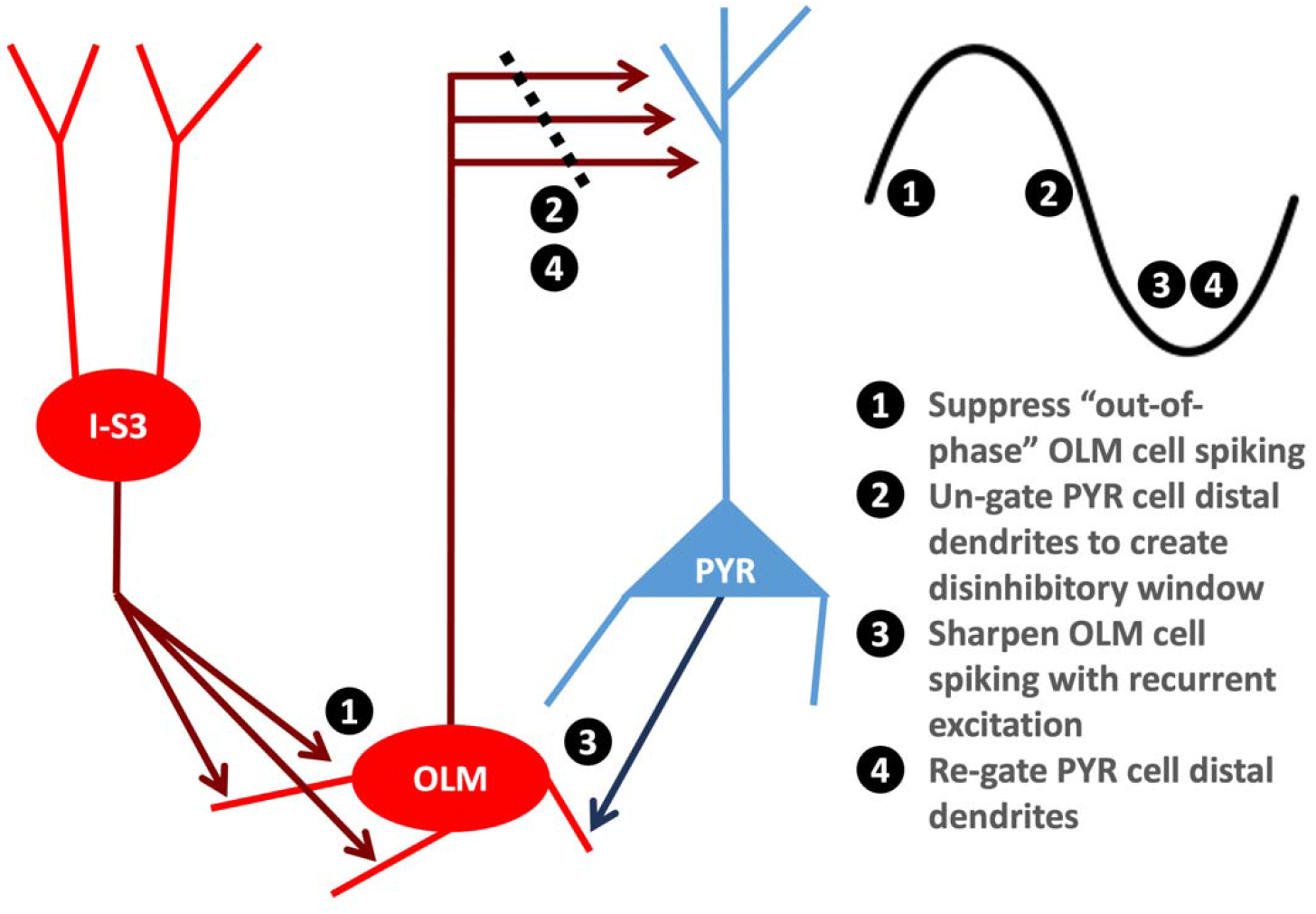
I-S3 cells are fit to suppress OLM cell spiking and disinhibit pyramidal cells. In our proposed mechanism, I-S3 cells control OLM cell spiking through spike suppression (step 1), leading to disinhibition of PYR cell distal dendrites (step 2), enabling excitation of OLM cells (step 3), and subsequent re-gating of PYR cell distal dendrites via OLM cell-mediated inhibition.

### Experimental Investigations

The predictions from our computational studies lead to several suggestions for experimental investigation. One precise prediction is that 30 I-S3 cell synapses were necessary to elicit inhibitory control over OLM cell spiking. If the variance-mean analysis estimation of three release sites by Tyan et al. (15) is correct, this would imply activation of at least 10 presynaptic I-S3 cells. Results from Tyan et al. (15) can also be re-tested by performing frequency-modulated optogenetic stimulation of CR+ cells while recording from OLM cells in the presence of excitatory synaptic blockers. If OLM cell frequency-modulated spiking control is then lost, the interpretation would be that the frequency modulation was due to feedforward disinhibition-driven excitation onto OLM cells. Additionally, instead of recording from OLM cells during these tests, one could record from PYR cells to verify frequencies at which I-S3 cell activation might lead to disinhibited spiking in PYR cells, if at all.

Another suggestion would be to perform closed-loop feedback stimulation with the theta-filtered LFP as has been done previously (21), but with photo-activation of CR+ cells at specific phases of theta. One could stimulate CR+ cells at phases closer to the peak versus the trough of theta, to assess whether the phasic timing of CR+ cells has an effect on the encoding and retrieval of information during a learning test such as the T-maze task. Our results predict an un-gating of pyramidal cells during theta phases that follow the activation of CR+ cells. On average (but with a large variance), I-S3 cells spike near the rising to peak phases of theta (16) which follows excitation from EC and precedes excitation from CA3. As such, if stimulated to spike at earlier phases, integration of inputs from EC by pyramidal cells could be favoured to promote sensory encoding. Likewise, if stimulated to spike at later phases, integration of inputs from CA3 by pyramidal cells could be favoured to promote retrieval of information.

Further, it is now clear that the location of OLM cells along the dorsoventral axis of the hippocampus matters in terms of whether one records high or low theta rhythms (22, 51, 52), possibly due to biophysical and connectivity differences with OLM cells in these different locations. As already noted above, modeling studies have suggested that whether h-channels are present or not in the dendrites of OLM cells could ‘control’ the theta frequency spike resonance. Here, we used OLM cell models with dendritic h-channels developed from intermediate CA1 mouse hippocampal data (27), and our PRC explorations indicated that current changes with inhibitory perturbations were mostly due to h-channels.

### Theoretical and Modeling Considerations

Our use of theoretical PRCs here is of a pulse-coupled type (53) where we used excitatory and inhibitory synaptic inputs to consider phase-locked entrainment of OLM cell models. PRCs could also be used to predict synchronous output, and for example, Type II PRCs imply synchronization with excitation (54, 55). We focused on the ability of our model OLM cells to exhibit phase advances or delays using inhibitory perturbations for I-S3 cell input considerations. Other studies have used synaptic perturbations with PRCs and with varying levels of neuron model complexity (36, 56).

Here, we have detailed multi-compartment models with a full suite of biophysical channel types, and since we wanted to situate our explorations in a realistic biological setting, our inhibitory perturbations took the form of dendritically distributed inhibitory synaptic inputs on the OLM cell models with synaptic features estimated from experiment. These PRCs serve as an approximation for how OLM cells would respond to repeated ‘periodic forcing’ (41), and they show that incoming inhibitory inputs mostly serve to slow down the firing of OLM cells. Moreover, by also examining the underlying changes in the biophysical currents of OLM cells, we found that h-channel changes dominate, but they are not sufficient to lead a phase advance with PIR mechanisms rather than a phase delay. Previous work has highlighted that T-type calcium and persistent sodium channels in deep cerebellar nucleus cell models can also underlie PIR spiking following long square negative current injections (57). In slight contrast, PIR spiking following long square negative current injections in our OLM cell models (**OSF Figure 1**) were initiated primarily by h-currents and then shaped by sodium currents, followed by T-type calcium currents, as well as several outward currents.

Another interesting aspect to consider are the more complex dynamical regime changes that occur *in vivo* during different awake and sleep cycles as well as behavioral states. While we have modelled *in vivo* states quite simply as high levels of AMPA-mediated synaptic bombardment, more complex neuromodulatory phenomena are likely to be governing state changes more closely associated with behavior. For example, experimental cholinergic modulation has previously been shown to switch the PRC in L2/3 pyramidal neurons from type II to type I (58), which would have synchronization consequences, and cholinergic inputs are known to be cell-specific (59). Other types of inputs to OLM cells have been reported previously, including serotonergic receptors (60), metabotropic glutamate receptors (61, 62), cholinergic receptors (63–65), additional nuances in NMDA/AMPA/Kainate receptors (66, 67), and TRPV1 receptors (68). However, in the absence of particular constraints, we opted to not include them at this time. It is unclear if a cholinergic-associated down-regulation of M-type current in our OLM cell models would enhance the small inhibitory synapse-driven phase advances that we report in our *in vitro* PRC analyses. However, given the phase invariance of non-h-type currents to inhibitory perturbations during rheobase spiking, we would not anticipate a large effect. Certainly, a more complex picture may exist in the *in vivo* state however.

To allow a focus on cellular details, we carried out virtual network explorations using detailed OLM cell models. Because we used models developed using an NMM approach, where a select number of neuronal model fits were closely matched with experimental recordings (27), we did not explore a full range of parameters to exhaustively assess the robustness of our findings. This can be done if we used the populations of ‘acceptable’ models that are generated for each optimization (27), but would require a considerable amount of computing resources to fully explore. The use of a virtual network allowed us to directly compare and translate *in vitro* aspects to *in vivo* to come up with our proposed specialized contribution of I-S3 cells via OLM cell control. Since *in vivo* recordings of specialized cell types is highly challenging, computational studies to develop hypotheses, predict and guide experimental studies are strongly needed. However, a virtual network is limited as it does not directly model all of the interacting network effects that can occur consequentially as a result of changes in the activation of the cell type of interest. At the same time, the large variety of neuron types and subclass interactions makes the consideration of interacting effects incredibly complex. Despite these interactions not being modelled explicitly, our approach using the detailed, biophysical OLM cell models offers the benefit of being able to dissect out putative mechanisms without having to build a full-blown circuit model initially, and has led to explicit suggestions for experimental investigation. Further, we note that recent computational (69, 70) and experimental (45) studies have begun to disentangle the co-activity dynamics of disinhibitory neurons with different neuron types in relation to behaviour. We would also like to note that our interpretations are in line with previous network modelling showing that VIP/SOM connectivity is sufficient for switching circuit activity between two processing modes where synaptic inputs in pyramidal cells are either integrated or suppressed (69, 71, 72).

While the use of virtual networks has limitations, building and using full microcircuit models requires other considerations with subsequent limitations. Two CA1 microcircuit models by Turi et al (49) and Bezaire et al. (39) have been built. The Turi et al model (49) consists of 150 cells that includes PYR cells and 6 types of interneurons including I-S3 cells. It is supportive of a disinhibitory role for I-S3 cells in encoding reward locations, and indeed, the disinhibitory effects of VIP+ cell activation on PYR cell spiking has been broadly reported in the literature (17). The Bezaire et al model (39) is a full-scale CA1 microcircuit model that consists of 338,740 cells including 8 types of interneurons, and shows the importance of some cell types over others in the production of theta rhythms. However, I-S3 cell types were not part of the Bezaire et al model. While both microcircuit models include OLM cells, the OLM cell models within them are not detailed in morphology (4 compartments) or biophysical currents relative to the developed OLM cell models used in the work here. Of particular note, no h-channels were included in the dendrites of the OLM cell models used in these microcircuit models. These are all reasonable limitations given the respective goals of these microcircuit models. In fact, there is an explosion of parameters at the circuit level (e.g., different cell types, connectivity parameters & strengths, network size, etc.) that usually necessitates a reduction in the resolution of analysis at the single cell level (e.g., dendritic compartments, ion channel types, etc.). A circuit model would also require a further expansion of high-performance computing resources towards simulating the full circuit model rather than focusing on the single-cell details and parameter spaces that we have focused on in this work. While our OLM cell models were developed carefully using an approach that directly matched the model development with the same biological cell (27), models at this level of detail do not currently exist for the many other interneuron types in the CA1. The huge endeavour of achieving this level of detailed models for many interneuron types in a full-scale CA1 microcircuit has been undertaken (73), although I-S3 cells have not yet been included.

### Summary & Conclusions

This work provides a deep understanding into how I-S3 cells could contribute to circuit function *in vivo*. This was accomplished by studying inputs to detailed OLM cells models and dissecting out how I-S3 cell inputs may contribute. By analyzing these factors carefully in biophysical computational models, we were able to generate predictions into the mechanisms through which I-S3 cells can effectively control OLM cell spiking and precisely coordinate modulation pathways. We also note that our interpretations are in line with previous network modelling showing that VIP/SOM connectivity is sufficient for switching circuit activity between two processing modes where synaptic inputs in pyramidal cells are either integrated or suppressed (69, 71, 72).

## Acknowledgements

This research was supported by the Natural Sciences and Engineering Research Council of Canada (NSERC): Discovery Grant (RGPIN -2016 -06182) to FKS and Graduate Scholarship (CGSD2 -504375 -2017) to AGM. We thank L. Topolnik for reading and providing feedback on this manuscript. We would also like to thank the reviewers for their various comments that helped us make significant improvements in the presentation of our work.

